# The geometry of hidden representations of protein language models

**DOI:** 10.1101/2022.10.24.513504

**Authors:** Lucrezia Valeriani, Francesca Cuturello, Alessio Ansuini, Alberto Cazzaniga

**Affiliations:** AREA Science Park, Padriciano 99, 34149, Trieste, Italy

## Abstract

Protein language models (pLMs) transform their input into a sequence of hidden representations whose geometric behavior changes across layers. Looking at fundamental geometric properties such as the intrinsic dimension and the neighbor composition of these representations, we observe that these changes highlight a pattern characterized by three distinct phases. This phenomenon emerges across many models trained on diverse datasets, thus revealing a general computational strategy learned by pLMs to reconstruct missing parts of the data. These analyses show the existence of low-dimensional maps that encode evolutionary and biological properties such as remote homology and structural information. Our geometric approach sets the foundations for future systematic attempts to understand the *space* of protein sequences with representation learning techniques.

## 1 Introduction

In the last years, deep learning models drastically changed the landscape of protein research, particularly for the prediction of structural and functional properties, giving a new impulse to technical and scientific advancements in this field. A particular class of deep learning models, which are referred to as protein language models (pLMs) [8, 20, 19, 16, 14], combine high predictive performance and architectural simplicity, making them an ideal candidate for evaluating hypotheses about computational strategies underlying their operations. pLM architectures have been heavily inspired by transformer-like models that emerged in the context of natural language processing: they consist of a stack of identical self-attention blocks trained in a self-supervised fashion by minimizing a masked language model (MLM) objective [24, 5]. It has been shown that the features learned by pLMs, after suitable fine-tuning, can be used to solve a wide range of supervised biological tasks [18, 26]; in this sense, these features possess some degree of universality. In these models, each module maps the data into a representation; it has already been observed that the organization of representations in the last hidden layer reflects biological and evolutionary information [8, 20], and the insurgence of similar properties in the attention matrices of the various blocks has been methodically investigated [25, 15]. In addition, analysis of other types of architectures highlighted that data representations in deep learning models undergo profound changes across the layers [2, 6]. Studying these behaviors is crucial for fully exploiting and understanding low-dimensional encoding of the data produced by the models. In this paper, we systematically investigate fundamental geometric properties of pLMs representations, such as their intrinsic dimension (ID) and neighbor composition, and find shared behaviors across many single-sequence language models trained by self-supervision on various protein datasets. Our main results are:

- Representations in single-sequence pLMs show a three-phased behavior revealed by global (ID) and local (neighborhood composition) measures
- The three phases, referring to the ID, consist of 1) a peak, in which the ID grows, reaches a maximum, and then contracts 2) a plateau, in which the ID is stationary across layers at low values and 3) a final ascent, in which the ID grows to return to values close to the one measured after the positional embedding
- Language models trained on multiple-sequence alignments (MSA) show a qualitatively different behavior. The ID stays approximately constant after a very feeble expansion at the first self-attention block
- In single-sequence pLMs, the neighbor composition of adjacent layers changes with a three-phased behavior tightly related to the ID one. After profound rearrangements in the peak phase, we find a plateau phase in which the neighbors’ relationships are approximately preserved followed by substantial changes again in the last phase
- The neighbor composition shows that evolutionary and structural information, such as remote homology and fold type, emerge gradually and then stabilizes across the layers in a way that is strictly consistent with the three-phased behavior.

Our findings shed light on the strategies employed by pLMs to solve the MLM task and can be a guide for effectively decoding the biological information distilled by their hidden representations, possibly also suggesting strategies to design more efficient, lightweight models.

## 2 Methods

The single-sequence pLMs we analyze are essentially characterized by the same architecture: after a learned positional encoding of the data, a stack of identical self-attention blocks transforms the input creating successive representations. These models are trained in a self-supervised way to perform a partial input reconstruction task by minimizing a masked language model loss. As a byproduct, the learned representations are rich in biological information. More in detail, the input data points, corresponding to proteins, are variable-length sequences of *l* letters *s* = *a*_1_,*a*_2_,…*a_l_*, chosen from an alphabet of *n_a_* (≃ 20) tokens corresponding to amino acids. Each token is encoded by an embedding layer into a vector of size *d*, so that the generic protein s is represented as a matrix *x*:= *f*_0_(*x*) of size *l* × *d*. A model with *B* blocks transforms a data point *x* ∈ ℝ^*l×d*^ into a sequence of representations: 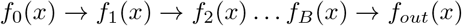, where *f_i_, i* = 1,…, *B* stands for the self-attention module at the i - *th* block, and the final LM-head *f_out_* is a learned projection onto dimension *l* × *n_a_*. The size of each hidden layer does not change across the model and is equal to *l* × *d*; therefore, the action of the model is a sequence of mappings ℝ^*l×d*^ → ℝ^*l×d*^. The representation of a protein across the network consists of a collection of *l* vectors that change across the layers, and several strategies for comparing variable length sequences have been investigated [4]. For each layer *i* we choose to perform global average pooling across the row dimension 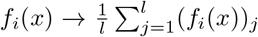, since this reduction retrieves sufficient biological information to solve, directly or possibly after fine-tuning, homology, structural and evolutionary tasks [8, 20]. For a given pLM, the action of the network on a dataset of *N* proteins can thus be described by *B* + 1 collections of *N* vectors in ℝ^*d*^: these will be the data representations that we will investigate. In our applications we will focus on representations obtained starting from ProteinNet [1] and SCOPe [10, 3], two biologically relevant benchmark protein datasets described in A.4. We will consider a selection of models from the ProtTrans and ESM families pre-trained on large protein databases, whose main properties are detailed in A.3. We also describe representations extracted from MSA-based pLMs in A.5.1.

### 2.1 Intrinsic dimension

The *manifold hypothesis* is based on the observation that many datasets embedded in high dimensions, resulting from the observations of natural phenomena (images, sounds, etc.), lie close to lowdimensional manifolds. The intrinsic dimension (ID) of a dataset is the dimensionality of the embedded manifold approximating the data; in other words, the ID is the minimum number of coordinates that allow specifying a data point approximately without information loss. We adopt the global estimator “TwoNN” of the ID developed in [9], which requires only local information on the distance to the first (*r*_1_) and second (*r*_2_) nearest neighbors of each data point, and that works under the mild assumption of approximately locally constant density. In such a case, the theoretical cumulative distribution *F* of the ratio *μ* = *r*_2_/*r*_1_ can be explicitly derived from the ground truth ID without information on the density; after approximating *F* with the empirical cumulate calculated on the dataset, one can estimate the intrinsic dimension. We refer to [9] for further details on the algorithm, and to A.5 for a description of the implementation adopted in our analysis of the ID in the hidden layers of pLMs. The “TwoNN” algorithm is robust to change of curvature and density, and it is asymptotically correct in the range in which the ground truth ID is ≈ 20. This estimator has already been employed to analyze representations in deep convolutional networks in [2].

### 2.2 Neighborhood overlap

The changes in data representation across the model can be traced in the rearrangement of the neighbor structure of the data space under the transformation induced by a block: points that are close in one layer may not be so in the following layer, and *viceversa*. The neighborhood overlap [6] measures the degree of similarity of two data representations by computing the common fraction of points that are *k*-nearest neighbors in both representations. Explicitly, consider the *k* points nearest to an element *x^i^* of the dataset at a given layer *l*, and let *A^l^* be the adjacency matrix with entries 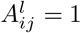 1 if *x^j^* is a neighbor of *x^i^* and 0 otherwise. The neighbor overlap between layers *l* and *m* is defined as 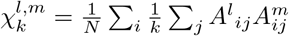, and it is easily seen to lie in [0,1]. The neighborhood overlap can be generalized in the following way. Let us consider a function *f* that associates a characteristic of interest to each data point. We can use f to define a neighborhood through the adjacency matrix 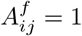 and 0 otherwise. In this case 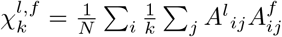 is the average fraction of neighbors of a given point in *l* that have the same property *f* as the central point. In [6] the authors consider the particular case in which *f* is a ground truth classification label. We focus on ground truth classes of biological interest such as protein fold, super-family, and family of the SCOPe dataset in 3.2.2 and A.6.4. In the case when *k* = 1, our measure collapses to the accuracy of first hit retrieval recently considered in [21].

## 3 Results

### 3.1 The intrinsic dimension has a characteristic shape for single-sequence language models

After each self-attention block, we extracted representations for proteins in ProteinNet, and plotted the ID against the block number, normalizing with respect to the total number of blocks (relative depth).

#### 3.1.1 The characteristic shape

The typical shape of the ID curve, that we found across diverse models trained on different datasets, has three distinct phases: a peak (**PE**) phase, a plateau (**PL**) phase and a final ascent (**FA**) phase (see Fig. 1, **A**). The peak develops early and occupies approximately the first third of the curve. In this phase, the ID rapidly expands, and after reaching a maximum in an ID range of a few tenths, it rapidly contracts. After achieving its maximum, the ID is compressed to remarkably low values that characterize the plateau, where the ID remains approximately stationary, reaching values of ≈ 6 - 7 at the elbow before the **FA**. In the **FA**, the ID grows again, going back progressively to values close to the ID computed on the representation after the positional embedding. The ID undergoes major changes across hidden layers: a ratio of ≈ 4 – 5 of the ID values can be observed between the minima at the **PL** phase and maxima at the **PE** phase. These changes are even more remarkable since the embedding dimension *d* remains unchanged across all the layers, depending only on the specifics of the single architectures reported in column “Emb. Dim.” of Table 1 in A.3.

**Figure 1:**
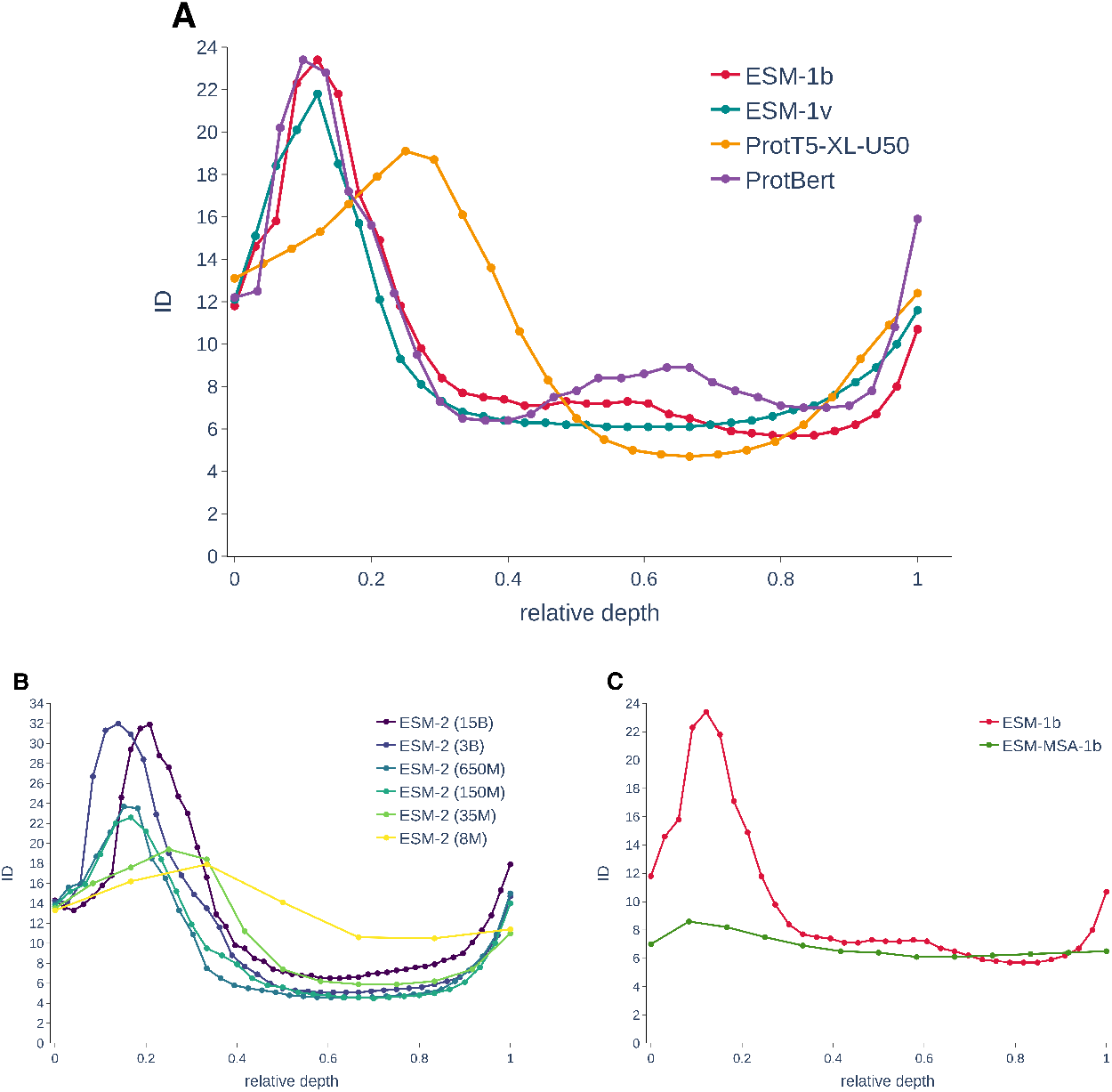
(**A**) ID in different models trained on different datasets show a three-phased behavior that consist in a peak (**PE**), plateau (**PL**) and a final ascent (**FA**). (**B**) The characteristic three phases in a sequence of models with common architecture and increasing size. (**C**) The ID in single-sequence (red line) and MSA-based pLMs (green line) are qualitatively different.

**Table 1:** Characteristics of the pLMs employed for extracting representations.

| Model | #Blocks | Emb. dim. | #Heads | #Params | Dataset | Reference |
| --- | --- | --- | --- | --- | --- | --- |
| ProtBert | 30 | 1024 | 16 | 420M | UR100 | [8] |
| ProtT5-XL-U50 | 24 | 1024 | 32 | 3B | UR50 \| BFD | [8] |
| ESM-1b | 33 | 1280 | 20 | 650M | UR50/D | [20] |
| ESM-1v | 33 | 1280 | 20 | 650M | UR90 | [16] |
| ESM-MSA-1b | 12 | 768 | 12 | 100M | UR50* | [19] |
| ESM-2(8M) | 6 | 320 | 20 | 8M | UR50/D | [14] |
| ESM-2(35M) | 12 | 480 | 20 | 35M | UR50/D | [14] |
| ESM-2(150M) | 30 | 640 | 20 | 150M | UR50/D | [14] |
| ESM-2(650M) | 33 | 1280 | 20 | 650M | UR50/D | [14] |
| ESM-2(3B) | 36 | 2560 | 40 | 3B | UR50/D | [14] |
| ESM-2(15B) | 48 | 5120 | 40 | 15B | UR50/D | [14] |

#### 3.1.2 Characteristic shape and model scale

We analyzed the effect of models size on the three-phased behavior described in 3.1.1 by computing the ID curve for the single-sequence model ESM-2 [14] in a range of size spanning more than three orders of magnitude, from 8M to 15B parameters. The three-phased behavior is preserved when changing the scale of the model (see Fig. 1, **B**), assuming that one considers sufficiently expressive architectures of size ≳ 35M. The maximum reached by the ID during the **PE** phase tends to increase accordingly to the scale of the models. In the plateau phase, we find a remarkable quantitative consensus on ID values which are approximately identical in models spanning two orders of magnitude in size, with number of parameters ranging from 35M to 3B, and with different values of extrinsic dimensionality *d*, ranging in values 480 — 5120. Increasing the size from 8M to 3B parameters, we progressively see the emergence of the typical shape. The largest model with 15B parameters stands out for the delayed peak location and the slightly higher ID value at the **PL** phase. Besides the size of the pLM, this may be related to the fact that ESM-2(15B) has been trained for roughly half the iterations with respect to the other models (see [14], Table S1, row “Training steps”).

#### 3.1.3 MSA Transformer has a different ID characteristic shape

MSA-based pLMs held a prominent role in the field [19]: they are the first deep learning model that largely outperformed direct coupling analysis [17] in unsupervised contact prediction, and they constitute a fundamental component in the Evoformer block of Alphafold2 [13] that revolutionized protein structure prediction. Despite the similar structure, MSA Transformer radically differs from single-sequence pLMs since the model is directly exposed to data containing rich homology information, and the self-attention block intertwines tight row attention and column attention layers, focusing on the co-evolution content of the MSA. These dissimilarities are reflected in the different behavior of the ID curve (see Fig. 1, **C**). The three-phased behavior disappears, replaced by a slight increase of the ID after the first self-attention block, followed by a slight decrease and a large plateau that extends for all the remaining layers. After the initial adjustment, the ID value of MSA Transformer representations (≈ 6 — 7) closely resembles the one observed at the **PL** phase in single-sequence models.

### 3.2 The neighborhood rearrangements mirror ID changes and provide evolutionary and structural information

#### 3.2.1 Neighborhood rearrangements are tightly related to the ID

We computed the neighborhood overlap (NO) of adjacent layers 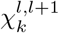 on the representations of ProteinNet considered in Fig. 1, **B**, where we analyze the ESM-2 model at different scales. The neighbor composition changes coherently with the three-phased behavior of the ID observed in Section 3.1.1, underlying an ID-NO relation not observed in the context of convolutional architectures [6]. The neighbors are subject to major rearrangements in the **PE** and **FA** phases, while they remain consistent in the plateau phase. In particular, for large enough models, size ≃ 150M, in the **PL** phase the neighborhood composition remains essentially unchanged, with ≈ 90% of shared neighbors across consecutive layers. The smallest model, characterized by low expressivity and high perplexity (see Table 2), constitutes an exception: major rearrangements can be observed throughout the layers, particularly in the plateau phase. We delegate further analysis on the NO of ESM-2 pLMs to Appendix A.6.2 and some considerations on the neighborhood composition of MSA Transformer to A.6.3.

**Table 2:**
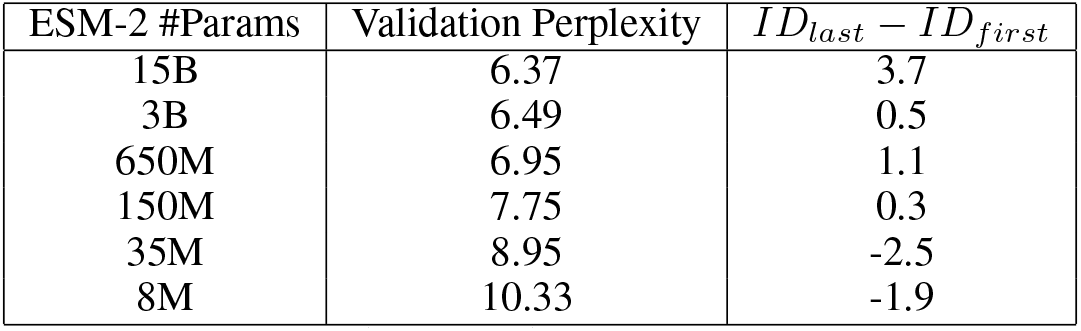
Validation perplexity [14, Table 1] and “reconstruction factor” calculated for ESM-2 at different scale.

| ESM-2 #Params | Validation Perplexity | $ID_{last} - ID_{first}$ |
| --- | --- | --- |
| 15B | 6.37 | 3.7 |
| 3B | 6.49 | 0.5 |
| 650M | 6.95 | 1.1 |
| 150M | 7.75 | 0.3 |
| 35M | 8.95 | -2.5 |
| 8M | 10.33 | -1.9 |

#### 3.2.2 Neighborhood structure and emergence of biological features: remote homology

Two proteins are said to be remote homologs if they correspond to highly dissimilar amino acid sequences while presenting a similar structure induced by common ancestry. It has been observed in [20] that Euclidean distances among representations in the last hidden layer of pLMs encode remote homology information. We study how this biological feature emerges in pLMs: considering representations of the SCOPe dataset, for every layer *l* we compute the neighborhood overlap 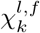 where *f* is the classification by super-family, excluding neighbors in the same family to focus on remote homology. Structural homology information is absent in the positional embedding layer; it grows smoothly in the **PE** phase reaching a stationary maximum 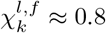 in the **PL** phase, and it suddenly decreases in the **FA** phase. We observe similar behavior for other biological tasks where f describes other types of homology relations in A.6.4, and we discuss in A.6.5 how this analysis could be beneficial for improving homology searches based on pLM representations [21].

## 4 Discussion and conclusions

In this work, we begin a systematic investigation of the geometric properties of representations in SOTA pLM with a twofold aim: 1) understand the computational strategies developed by the models to solve the MLM task and 2) monitor how evolutionary and structural information develops across layers. We observe a three-phased behavior shared across diverse single-sequence pLM models. These phases are clearly outlined by the variation of ID that shows a peak, a plateau, and a final ascent (see Fig. 1). The ID changes but remains orders of magnitude smaller than the size of the hidden layers, and the constant size of the layers rules out the potential confounding factor of variable layer size in the pLMs we consider. We conjecture that the initial ID expansion favors the creation of linearly separable features that can be more easily combined and processed downstream, mimicking the dimensionality expansion characteristic of kernel methods [11]. In this phase, the neighbors rearrangements reflect the progressive emergence of evolutionary and structural information (Fig. 2, **B**). The plateau is characterized by the presence of low-dimensional representations ID ≈ 6 - 7 in which evolutionary and structural information is maximal (Fig. 2, **B**). In the final ascent, the ID grows probably because the representation has to be enriched with less abstract features that better support the complex decision-making necessary to perform well on the reconstruction task. Accordingly, the neighbor structure reflects less explicitly the evolutionary and structural information. This picture is consistent across models spanning orders of magnitude in complexity and size or representations (Fig. 1, **B**). The bigger the model, the higher the ID peak, hinting at a larger diversity and a better separability of the features extracted, which goes along with the lower perplexity. The low ID values that characterize the plateau phase are largely independent of the scale. MSA-based pLMs present an approximately constant ID after a very feeble expansion at the first self-attention block. The visible difference between single-sequence and MSA-based pLM is possibly due to the fact that single-sequence models are in charge of discovering homology relationships that in MSA Transformer are provided as an input. As a final remark, we observe that, while the ID estimator we adopted is a global measure of dimensionality, the neighbor structure is inherently local. Remarkably, these properties change together in perfect unison, and we left a more careful investigation of this phenomenon as an open question for future research, as well as a deeper dive into the probabilistic structure of these representations [7].

**Figure 2:**
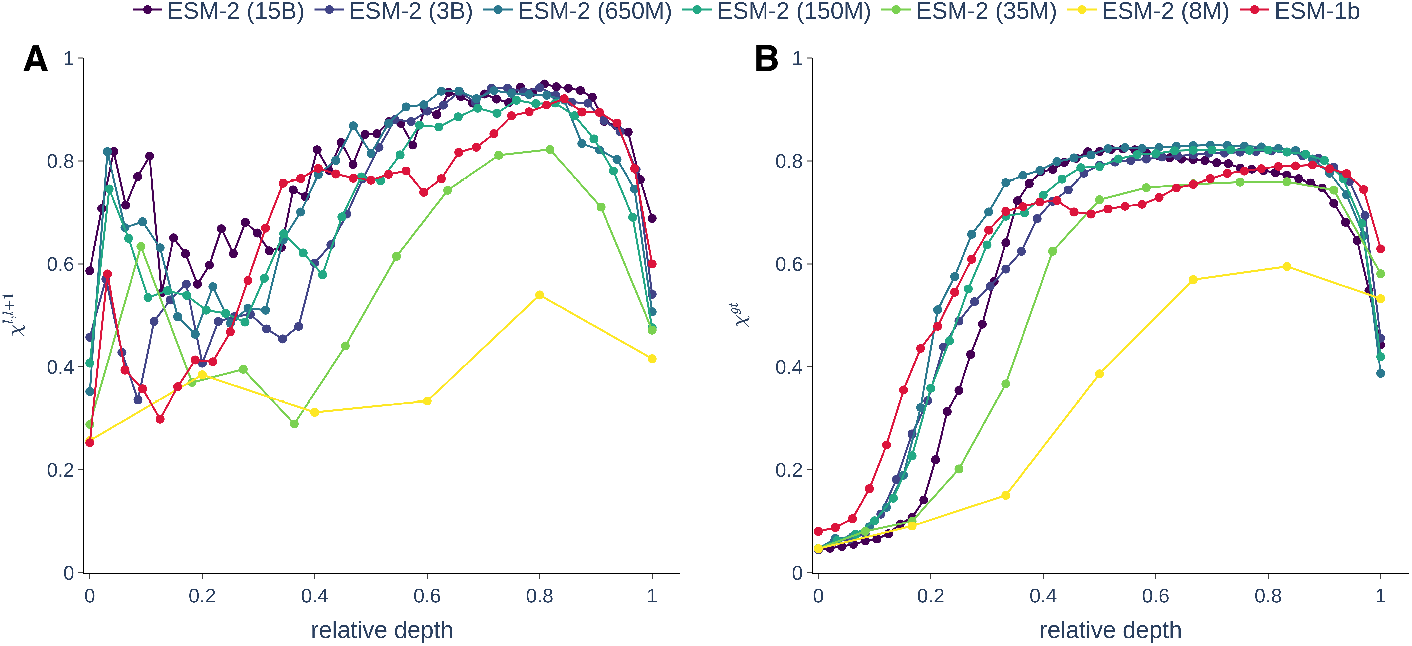
(**A**) The neighborhood overlap computed for consecutive layers shows a major restructuring of neighbor relationships in the **PE** and **FA** phase and minor changes in the **PL** phase, thus confirming the three-phased scenario inferred from the ID analysis.(**B**) Remote homology information is gradually acquired during the **PE** phase, it remains stationary in the **PL** phase, and is then partially lost in the **FA** phase.

## Acknowledgments and Disclosure of Funding

The authors acknowledge AREA Science Park supercomputing platform ORFEO made available for conducting the research reported in this paper, and the technical support of the staff of the Laboratory of Data Engineering. We thank Alessandro Laio and Ginevra Carbone for feedback on the manuscript and insightful conversations. L.V. was supported by the grant BOL “BIO Open Lab”. F.C. was supported by the grant PNR “FAIR-by-design”. A.A. and A.C. were supported by the ARGO funding program.

## A Appendix

### A.1 Experimental setup

**Hardware** All experiments were performed on a machine with 2 Intel(R) Xeon(R) Gold 6226 with a total of 48 threads, 256GB RAM equipped with 2 Nvidia V100 GPUs with 32GB memory, hosted on the ORFEO supercomputing platform at AREA Science Park. The GPUs were used to generate embeddings and to compute nearest neighbors.

### A.2 GPU kNN search

The nearest neighbor searches for the calculation of the neighborhood overlap were carried out by means of the python interface of the Facebook AI Similarity Search library [12], version 1.7.2. The library is particularly suited for large datasets embedded in high dimensions, since it is based on a reliable approximate and extremely fast similarity search procedure.

### A.3 Models

We consider two sets of transformer-based protein language models: ProtTrans [8] and Evolutionary Scale Modeling (ESM) [20, 19, 16, 14]. In our analysis, we consider all the models listed in Table 1, where we report the number of self-attention blocks, the embedding dimension, the number of attention heads, the total number of parameters, and the dataset used for pre-training the model. For further details on the architectures of the pLMs and the self-supervised training procedure, we refer to the original reference reported in the last column of Table 1.

### A.4 Datasets

#### A.4.1 The ProteinNet dataset

ProteinNet [1] is a standardized dataset for evaluating protein sequence-structure relationships recommended in [18] for assessing the contact prediction task. We use the ProteinNet training set, composed of 25299 sequences, as a reference for extracting the pLM representations used to analyze the characteristic ID curves in Fig. 1. The same representations were used to compute the neighborhood overlap of consecutive layers in Fig. 2, **A** and for the analysis of Fig. 4.

**Figure 3:**
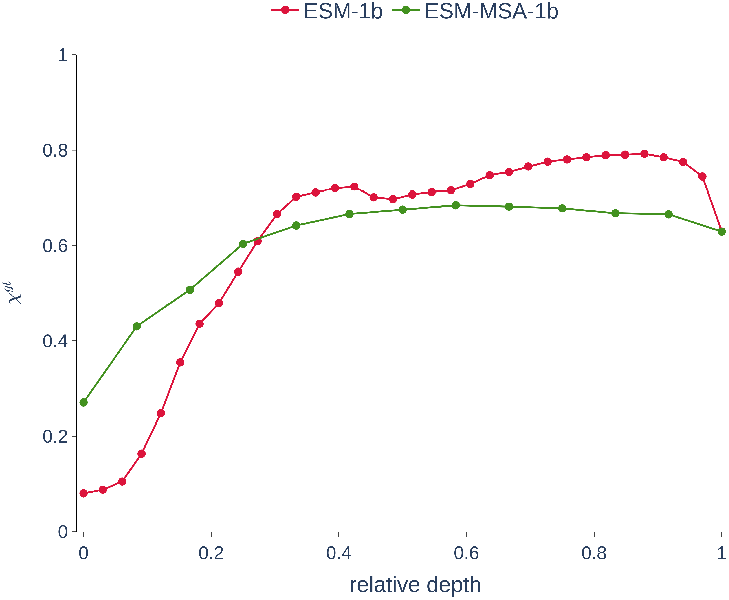
Remote homology information, given by SCOPe super-family classification, is present in the positional embedding layer of MSA Transformer (green curve), and it rapidly saturates to a value of ≈ 0.6.

**Figure 4:**
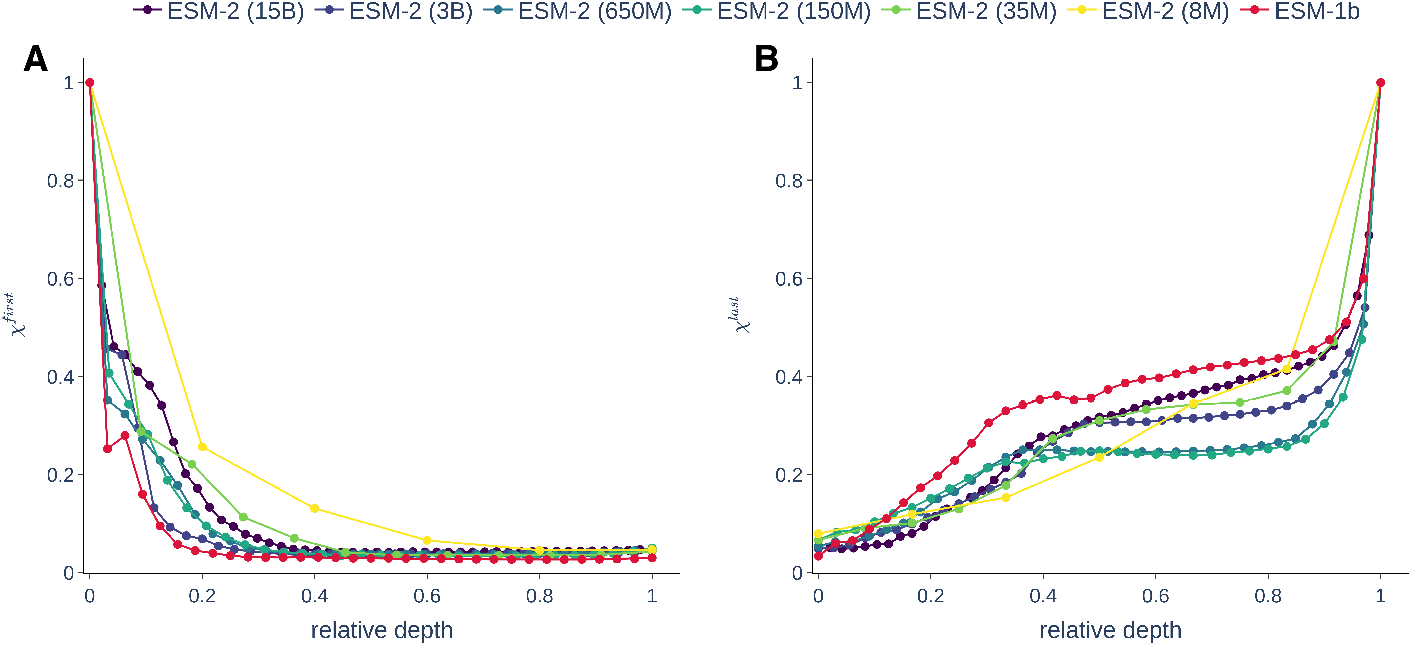
(**A**) Neighborhood overlap computed with the positional embedding layer (“first”) shows that similarity with the embedded input is lost after the **PE** phase. (**B**) Neighborhood overlap with the last hidden layer (“last”) shows that similarity with the representation used for solving the MLM task emerges feebly during the **PL** phase and increases rapidly during the **FA** phase.

#### A.4.2 The ProteinNet-MSA dataset

For each sequence in the ProteinNet training set, we obtain an MSA by following the procedure reported in [19, Section A.2], which is based on a search of the UniClust30 dataset with HHblits [23]. The MSA obtained following this recipe are then filtered using HHfilter [23], setting `-n 256`. This procedure, introduced in [19], maximizes sequence diversity while ensuring that the MSA size satisfies the dimensional constraints, allowing ESM-MSA-1b inference to take place in GPU. Thus, we obtain the MSA-ProteinNet dataset of 25299 MSAs with a maximum depth of 256. Each dataset element contains a protein in the ProteinNet as the first sequence in the alignment.

#### A.4.3 The SCOPe dataset

For our analyses of neighborhood overlap devoted to understanding the identification of remote homology relations in hidden layers, we consider the Astral SCOPe v2.08 dataset [3], containing genetic domain sequence subsets filtered to obtain < 40% pairwise sequence identity. Each domain is hierarchically classified into fold, super-family, and family. We impose an initial filter by excluding the Rossman-like folds (c.2–c.5, c.27 and 28, c.30 and 31) and the four- to eight-bladed b-propellers (b.66–b.70), as recommended in [22]. As a result, we obtained a dataset composed of 14535 sequences. For each specific task, we define an ad hoc dataset, which we will describe in detail in the corresponding sections A.5.2 and A.6.4, essentially to ensure sufficient population in the classes.

### A.5 Experiments

#### A.5.1 Intrinsic Dimension characteristic curve

**Two Nearest Neighbors ID estimator** To estimate the intrinsic dimension of hidden representations, we use the Two-Nearest Neighbors-Based (“Two-NN”) ID estimator [9]. The algorithm is based on a simple analytical result: under the hypothesis of a uniform density of points in ℝ^*d*^, the cumulative probability distribution of the random variable 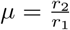, where *r*_1_, *r*_2_ are respectively the distance to the first and the second neighbor of a given point, is given by *F*(*μ*) = 1 - *μ*^−*d*^. Therefore, for a given dataset whose points are indexed by *i* = 1,…, *N* in ℝ^*D*^ (with *D* >> *d* in interesting cases), we compute for each point the ratios *μ_i_*, sort them in ascending order with a permutation *σ*, and, by defining the empirical cumulative distribution 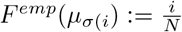, we can obtain an estimate of d as the slope given by a linear regression (passing through the origin) of the following variables: 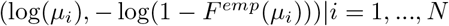. The Two-NN algorithm requires minimal information: the distances to each point’s first and second nearest neighbor; therefore, the strong hypothesis of a uniform density used to obtain the main result can be relaxed to a weak assumption of *local* uniformity. We use the code in [2] https://github.com/ansuini/IntrinsicDimDeep to estimate the ID and its reliability through a progressive, random decimation process that allows testing the stability of the result with respect to a change in spatial scale. Since the estimate is approximately scale-invariant, we take the ID estimate as the mean over the values collected during the decimation.

**Single-sequence representations** For extracting representations from single-sequence protein language models, we follow the recommendations reported in the repository of ESM (https://github.com/facebookresearch/esm) and ProtTrans (https://github.com/agemagician/ProtTrans). After extracting the representations, we apply global average pooling along the row dimension (that keeps track of the position in the sequence) as described in Section 2.

**MSA Transformer representations** We describe the procedure for extracting representations from an MSA-based pLM in more detail since it differs from the one described in Section 2. An element *q* in the MSA-ProteinNet dataset A.4.2 consists of a matrix of *m × l* tokens corresponding to an amino acid identity or a gap introduced aligning to the query sequence *s*, which is placed in the top row of *q*. We perform inference via ESM-MSA-1b, collecting hidden representations corresponding to the positional embedding *x*:= *f*_0_(*x*) and to the result of the *B* = 12 transformations *f_i_*(*x*) of self-attention blocks, which intertwines row and column attention layers. Each representation is a tensor of dimension *m × l* × *d*, where *m* is the depth of the MSA, *l* is the length of the ProteinNet query sequence on top of the MSA, and *d* = 768 is the embedding dimension. Following the main practice in the literature, we contract the first two indices of the tensor obtaining a *d* dimensional vector 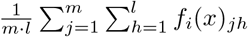. Thus, to each sequence in ProteinNet, we associate *B* + 1 vectors of dimension d and, repeating the operation on all *N* = 25299 proteins, we obtain the *B* + 1 representations of ProteinNet whose geometric structure is the focus of our ID estimation and analysis in Section 3.1.3.

#### A.5.2 Neighborhood overlap

**Remote homology detection in hidden layers: super-family belonging and neighborhood overlap** In our study on the overlap of the neighborhood structure of layer representations and the classification with respect to a remote homology task, we select proteins in the SCOPe dataset belonging to superfamilies with at least ten elements. Furthermore, we ensure that each super-family is composed of at least two families. We thus obtain a dataset composed of 10256 sequences grouped in 288 super-families. At evaluation time, when computing the *k*-NN of a given protein domain we remove elements in the same family to ensure we are considering only sufficiently remote homologs. We set *k* = 10 throughout our experiments, but it can be shown that the neighborhood overlap behavior is qualitatively conserved varying the number of neighbors *k*.

We repeat the same experiment on super-family neighborhood overlap for representations of MSA Transformer and compare it to the single-sequence model ESM-1b in Fig. 3. The behavior of the MSA Transformer neighborhood overlap curve (green curve) is qualitatively and quantitatively different from the single-sequence one (red curve): 1) part of the remote homology information is already present in the positional embedding layer; 2) the neighborhood overlap increases steeply in the first three blocks; 3) the level of agreement of neighborhood structure and super-family belonging saturates and remains approximately constant after the fourth block.

### A.6 Other experiments

#### A.6.1 Final ID expansion is related to perplexity

Perplexity is a measure of uncertainty of a language model in predicting masked elements of a sequence [14, Supplementary 1.1.2]. In particular, in [14] the authors show that the perplexity of an ESM-2 model, calculated on a validation set, is highly correlated with its structure prediction performance on CASP14 and CAMEO test sets. We focus instead on the “reconstruction factor” of an ESM-2 model, measured as the difference of ID measured at the “last” and at the “first” hidden layer. Validation perplexity and “reconstruction factor” are anti-correlated, with a Pearson correlation coefficient of —0.841 at p-value 0.036 computed from the corresponding columns in Table 2.

#### A.6.2 Neighborhood rearrangement in first and last hidden layer

In analogy with Section 3.2.1, we study the evolution of the neighborhood structure along a pLM measuring the neighborhood overlap between representations of the layer after positional embedding (“first”) and a generic layer l (see Fig. 4, **A**). The local structure of representations after the embedding of the input is entirely rearranged once the **PL** phase is reached.

Similarly, we calculate the neighborhood overlap between representations of the layer after the last self-attention block (“last”) and a generic layer l (see Fig. 4, **B**). The neighborhood organization of the representations at the “last” layer emerges gradually reaching a value of ≈ 0.3 at the **PL** phase. The neighborhood overlap increases steeply during the **FA** phase.

#### A.6.3 Neighborhood rearrangement in MSA Transformer

We study the neighbor rearrangement in the different layers of MSA Transformer and compare it to ESM-1b to investigate the impact of exposure to evolutionary data on the local geometry of representations. Results in Fig. 5, panel **A**, show that MSA Transformer retains some of the initial neighborhood structure in all layers. In contrast, the information of the embedding layer is entirely rearranged in ESM-1b representations. Neighborhood overlap of MSA Transformer and ESM-1b representations with the respective “last” hidden representations have similar qualitative behavior (see Fig. 5, **B**). However, the consensus with the “last” hidden representation is reached faster in MSA Transformer that present a high neighborhood overlap of ≈ 0.6 already at the **PL** phase. After the first self-attention block, MSA Transformer successive layers share most of the neighborhood structure (see Fig. 5, **C**). In particular, with respect to ESM-1b representations, we observe a lower level of rearrangement in the **PE** phase and an area with a high neighborhood overlap of successive layers that extends longer than the **PL** of single-sequence models.

**Figure 5:**
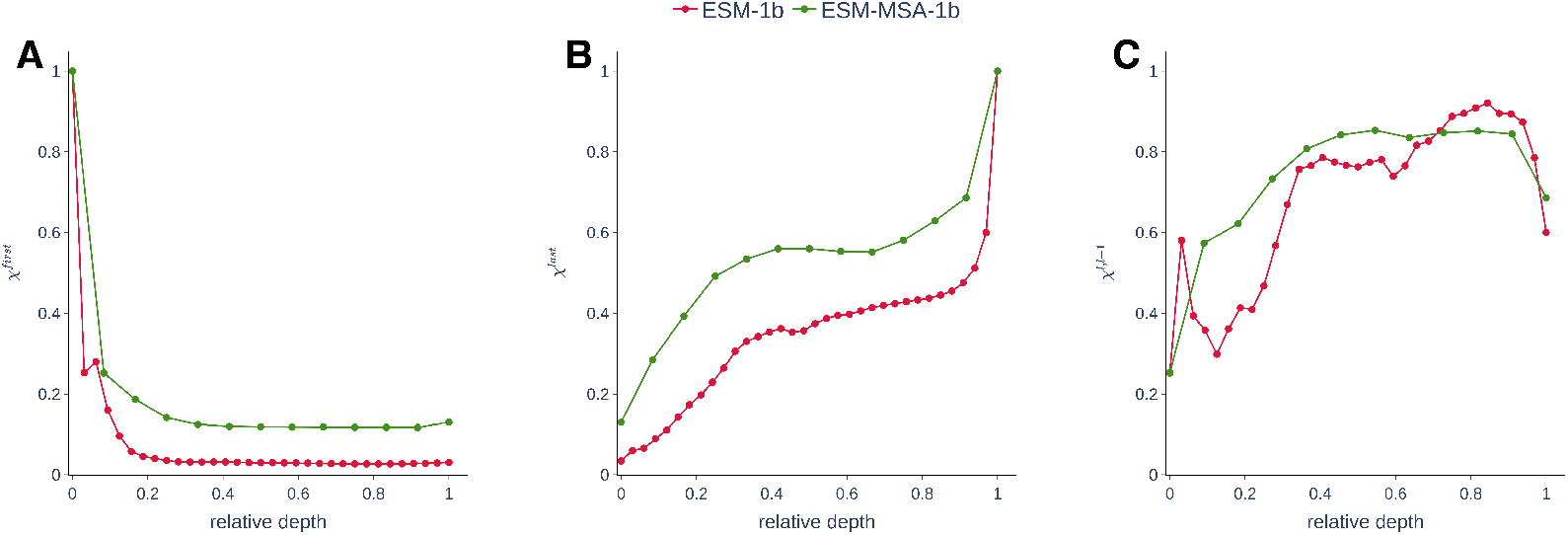
Neighborhood overlap with respect to the layer after positional embedding (“first”, panel **A**), to the layer after the last self-attention block (“last”, panel **B**) and values (panel **C**), for ESM-MSA-1b (green) and the ESM-1b model (red).

#### A.6.4 Homology task: emergence of fold, super-family and family relations in hidden layers

We study the emergence of homology relations in the geometric organization of hidden layers representations of single-sequence pLMs of the ESM family, studying neighborhood overlap with fold, super-family, and family classes of the SCOPe dataset. We select only proteins in fold, superfamily, and family classes of SCOPe containing at least 20 sequences, and we set the number of neighborhoods *k* = 10. We obtain three datasets, one for each class: the fold dataset contains 10926 domain sequences classified in 165 folds, the super-family has 8580 entries classified in 167 super-families, and the family dataset is made of 3610 sequences belonging to 91 families.

The results reported in Fig. 6 point out a general trend similar to that observed for the remote homology task of Section 3.2.2. After a gradual increase in the **PE** phase, the neighborhood structure aligns well with separation in classes in the **PL** phase and decreases suddenly in the **FA** phase. The maximum neighborhood overlap is reached at the **PL** phase for all the homology tasks, and it exceeds 0.7 for models with more than 35M parameters, showing that various types of homology relations are best encoded in the low-dimensional representations at the plateau.

**Figure 6:**
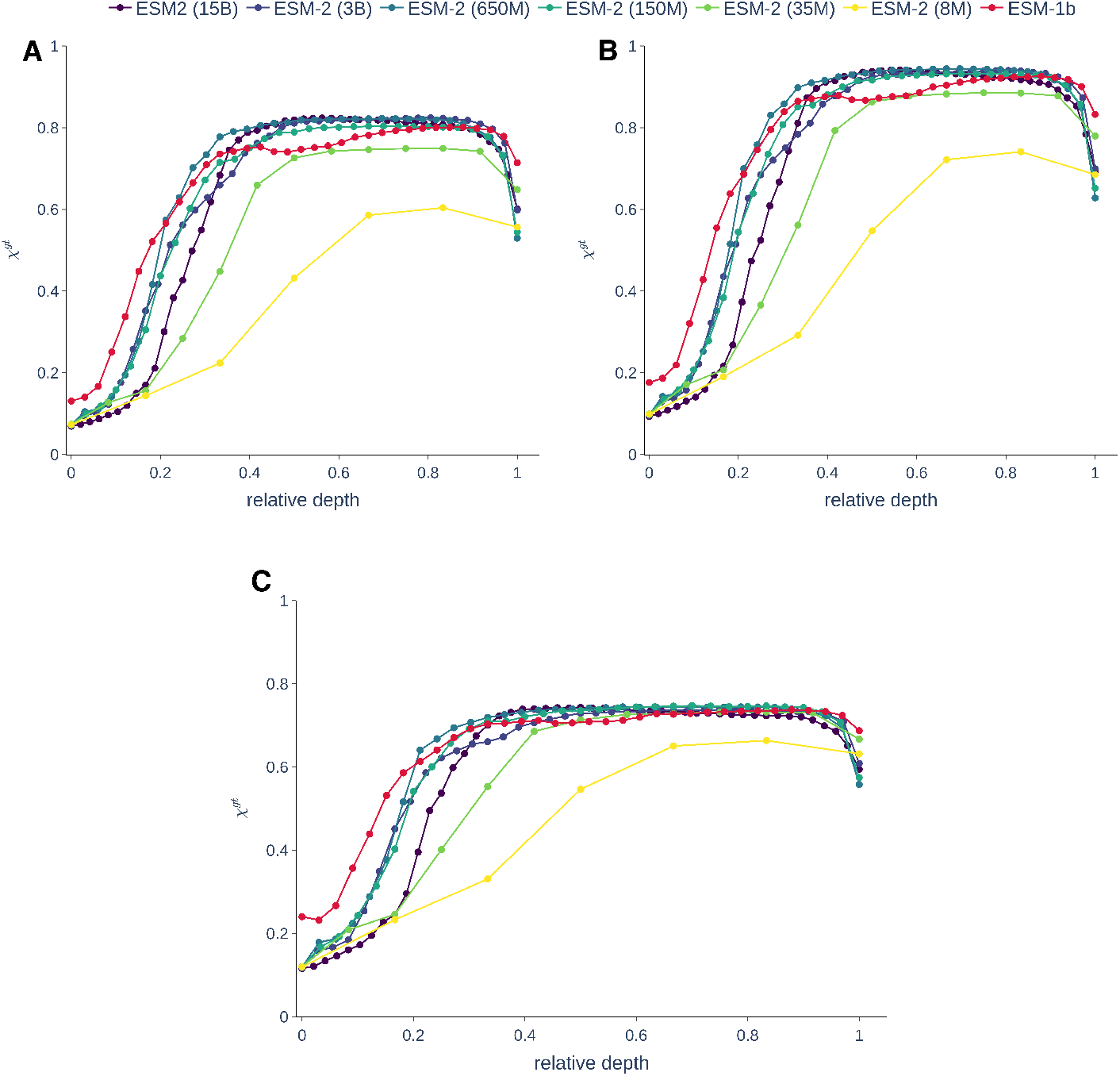
Neighborhood overlap with respect to fold (**A**), super-family (**B**), and family (**C**) ground truth classes for ESM single-sequence models.

#### A.6.5 Nearest neighbor search in PL layers improves identification of protein relations

It was recently shown in [21] that first nearest neighbor searches for remote homologous protein domains based on the last hidden layer representations of large pLMs outperforms SOTA methods based on sequence similarity. Adapting the approach in 3.2.2, we mimic the experiment performed in [21, Section 2] by 1) considering protein domains in SCOPe belonging to a super-family with at least 2 sequences, 2) setting the number of neighbors to *k* = 1.

From the results in Fig. 7 it emerges that, for the ESM-1b and the ProtT5-XL-U50 models, considering representations in the **PL** layer improves the accuracy of the 1-kNN homology search. In particular, for ProtT5-XL-U50 we have an improvement of ≈ 6% performing the search on a **PL** layer instead of in the last layer before the output. It will be interesting to assess if an analogous behavior is confirmed in experiments reproducing the exact conditions in [21, Section 2].

**Figure 7:**
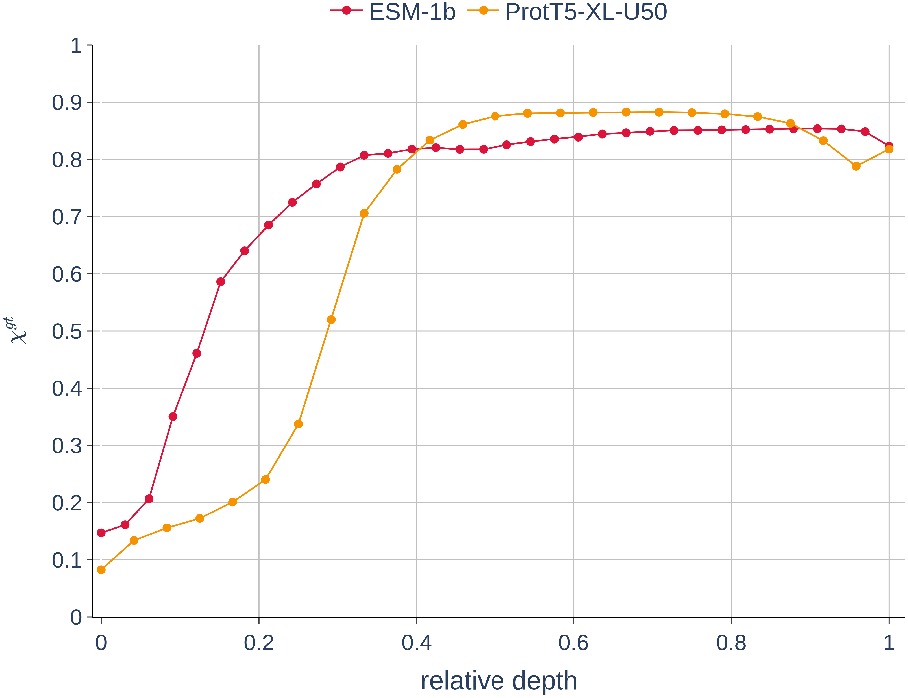
First nearest neighbor SCOPe super-family retrieval accuracy of ESM-1b and Prot-T5-XL-U50 is higher in **PL** layers.

